# How fluorescent tags modify oligomer size distributions of the Alzheimer-peptide A*β*(1-40)

**DOI:** 10.1101/372136

**Authors:** J. Wägele, S. De Sio, B. Voigt, J. Balbach, M. Ott

## Abstract

Within the complex aggregation process of A*β*-peptides into fibrils, oligomeric species, play a central role and reveal fundamental properties of the underlying mechanism of aggregation. In particular, low molecular weight aggregates have attracted increasing interest because of their role in cytotoxicity and neuronal apoptosis, typical of aggregation related diseases. One of the main techniques used to characterize such early stages of aggregation is fluorescence spectroscopy. To this end, A*β*-peptide chains are functionalized with fluorescent tags, often covalently bound to the disordered N-terminus region of the peptide, with the assumption that functionalization and presence of the fluorophore will not modify the process of self-assembly nor the final fibrillar structure. Up to date, experimental findings reveal size distributions of thermodynamically stable oligomers ranging from very narrow distributions of dimers to octamers, to very broad distributions up to 50-mers. In the present investigation we systematically study the effects of five of the most commonly used fluorophores on the aggregation of A*β*(1-40)-peptides. Time-resolved and single-molecule fluorescence spectroscopy have been chosen to monitor the oligomer populations at different fibrillation times, TEM, AFM and X-ray diffraction to investigate the structure of mature fibrils. While the structures of the mature fibrils were only slightly affected by the fluorescent tags, the sizes of the detected oligomeric species varied significantly depending on the chosen fluorophore. In particular, we relate the presence of high molecular weight oligomers (as found for the fluorophores HiLyte 647, Atto 647N and Atto 655) to net-attractive, hydrophobic fluorophore-peptide interactions, which are weak in the case of HiLyte 488, and Atto 488. The latter form low molecular weight oligomers only. Our findings reveal the potentially high impact of the properties of fluorophores on transient aggregates which needs to be included in the interpretation of experimental data of oligomers of fluorescently labeled peptides.

## INTRODUCTION

Amyloid forming proteins have attracted a great deal of attention in the last decades because of their association with a numerous group of degenerative conditions. Alzheimer’s, Huntington’s, Parkinson’s, Creutzfeldt-Jacob and prion diseases are some examples of the most well-known neuro-degenerative disorders, while the most well-known systemic ones are amyotrophic lateral sclerosis and type II diabetes (1). The etiology of all the aforementioned diseases seems to be found in the "defective" folding or "misfolding" of normally soluble, functional peptides and proteins, and their subsequent conversion into intractable aggregates, also known as amyloid fibrils (2). The latter, however, have been found to be a well-defined structural form also for many proteins unrelated to diseases (3). This observation suggests, in fact, that the structural motif common to these aggregates is broadly accessible by various polypeptide chains and can also be functional (4, 5). Either way, amyloidogenic proteins seem to follow a common dynamical pathway towards fibrils formation, characterized by an intermediate step of aggregation into small heterogeneous oligomers. The latter are also deemed to be the primary cause of cyto- and neurotoxicity (6). In Alzheimer’s disease, for instance, recent studies have demonstrated that cytotoxicity is indeed associated with the smaller still soluble species of Aβ oligomers, which lead to cell death by binding to the neuronal membrane and possibly by membrane permeabilization (7–11). The two most abundant amyloid allomorphs in the extracellular aggregates signature of advanced Alzheimer’s disease are A*β*(1-40) and A*β*(1-42) (12, 13). Generated by *β*-secretase cleavage of the intra-membrane amyloid precursor protein (APP) (14), the latter peptides have an intrinsically disordered structure which can be subject to conformational changes towards secondary and tertiary structures. Numerous *in vitro* studies of short and medium length amyloidogenic peptides have provided some clues to amyloid formation with an α-helix to *β*-sheet folding transition. However, the mechanisms triggering such conformational changes as well as aggregation are still unclear both *in vivo* and *in vitro* (1). Efforts to develop a more mechanistic understanding of how A*β* assembles into toxic species have been limited by significant experimental challenges. To start with, A*β* normally circulates in plasma and cerebro-spinal fluid as a soluble peptide in nanomolar to picomolar concentrations (2, 15, 16) and its aggregates are metastable and highly heterogeneous (17–19). Additionally, the self-assembly dynamics are highly dependent on the environmental conditions like pH, temperature, ionic strength or presence of metal ions (20) as well as concentration (21). Each of those factors can have enhancing or impeding effects in respect to aggregation, thus protocols on how to handle such peptides should be selected with great care, trying to keep conditions as close as possible to the physiological ones. Given the especially low concentrations at which A*β* peptides are normally found, many of the most popular techniques, such as polyacrylamide gel electrophoresis (PAGE) or size exclusion chromatography (SEC), are not suitable for these studies. Thus, investigation methods which can work at much lower concentrations, as optical techniques based on absorption and/or fluorescence detection, have been most widely employed to monitor in real-time the kinetics of the early aggregation processes in solution (20). In the last two decades techniques as fluorescence absorbance (22), photobleaching (19, 23), self-fluorescent-quenching (20), fluorescence correlation spectroscopy (FCS) (24, 25), fluorescence cross-correlation spectroscopy (FCCS) (26), dual-color fluorescence cross-correlation spectroscopy (dc-FCCS) (27), Förster resonance energy transfer combined with fluorescence correlation spectroscopy (FRET-FCS) (26), confocal two-color coincidence detection (cTCCD) (28), but also fluorescence imaging of labeled peptides (19, 29–31) or binding of Thioflavine T (ThT) (32) or Congo Red (33), have been widely applied for the study of such systems at very low concentrations. Despite the great sensitiveness of such optical techniques, one draw-back about using fluorescence is to be found in the inherent, yet necessary, modification of the original peptide system due to the attached fluorophore. The latter modification happens either through binding to aggregate-selective fluorophores, like Congo-Red or ThT, or through N-terminus covalent binding (see Fig. 1A). The N-terminus is considered to be a loose end of the primary peptide structure, which is thus supposed not to strongly contribute to its aggregation mechanisms nor structure formation (19, 28, 30). Nonetheless other experimental studies have suggested quite the opposite. Modifications of the N-termius (34, 35) or labeling with various fluorophores (36) can, in fact, interfere and modify the A*β* peptide self-assembly dynamics and as a result, its structure formation. In support to these findings, more recent theoretical and numerical studies have also pointed out that chemical modifications of the protein surface, as the addition of a fluorophore, do affect the physical properties of protein solutions. In particular, the chemical modification is found to be equivalent to the addition of a large hydrophobic patch with a large attractive potential energy well, significant even at low labeling fractions (37). In order to experimentally address the effect of N-terminus bound fluorophores, we studied the early stages of aggregation of fluorescently labeled A*β*(1-40) with five different fluorophores by time-resolve and single-molecule fluorescence experiments. More in detail, we have investigated the following fluorophores: Atto488, Atto647N and Atto655 and HiLyte488 and HiLyte647 by a concerted approach of different techniques. With the help of time-resolved and single-molecule fluorescence spectroscopy we could carefully monitor and characterize the early stages of aggregation of all the above listed samples at a single molecule level. X-ray diffraction measurements have been used to test the local structure of the final fibrils while imaging techniques such as atomic force microscopy (AFM) and transmission electron microscopy (TEM) have been used to directly visualize eventual effects translated up to the mesoscopic scale, such as changes in length or thickness, undetectable otherwise. As a result from the single molecule studies we found different characteristic distributions of oligomer sizes which we relate to specific properties of the fluorophores. Fluorophores displaying net-attractive interactions with the peptide, lead to stable high molecular weight (HMW) oligomers, which are not further participating in the fibril formation. On the other hand, fluorophores displaying less hydrophobic interactions, tend to form low molecular weight (LMW) oligomers which are consumed with ongoing fibrillation. From a structural point of view, TEM and AFM images revealed a diverse scenario of mature fibrillar structures ranging from very straight and thick to thin and curvy, while the local structure of the *β*-sheets investigated via XRD showed no notable differences between differently labeled A*β*(1-40)-peptides. All the observations collected thus far encouraged the conclusion that nucleation and growth are restricted to small (labeled) oligomers without structural deviations from the wild type oligomer: (i) XRD measurements show no sensible variation in the structure of the nucleation units nor (ii) is the fibrillation kinetics changed; (iii) the detected stable HMW aggregates do not contribute to fibril formation as they stay detectable by single molecule fluorescence and visible in AFM. Structural differences are only detected on the macroscopic scale, as supported by TEM images.

**Figure 1:**
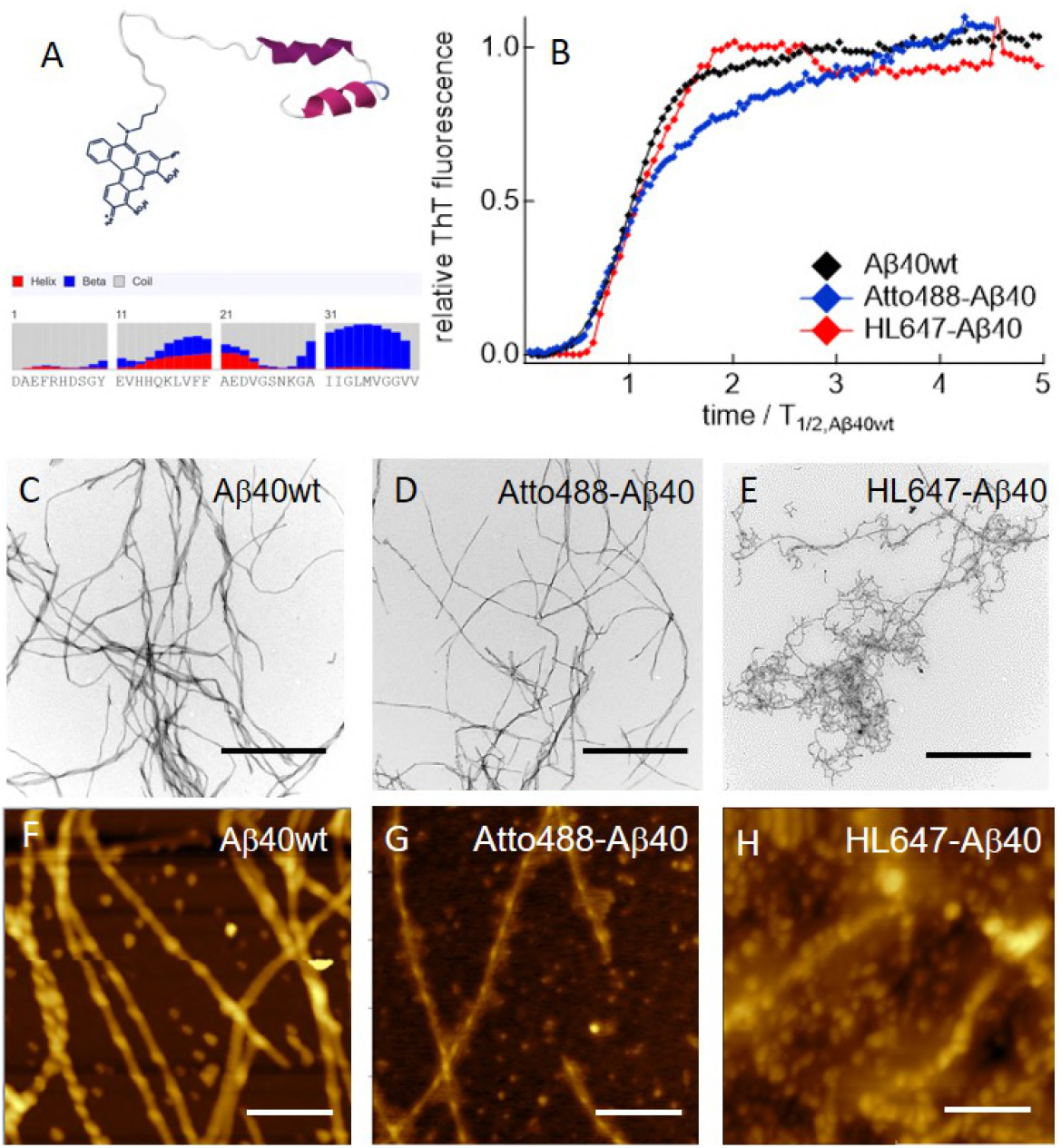
The effect of N-terminus bound fluorophores on the fibrillation kinetics and fibril morphology of A*β*40. A: Fluorescence labeling of the unstructured N-terminus of A*β*40 is expected to have the least effect to structure formation (structure model: PDB 2lp1). Inset: amino acid sequence of A*β*40 with secondary structure prediction using RaptorX (39). B: The time dependent ThT fluorescence of A*β*40wt without (black) and with 5% of Atto488-A*β*40 (blue), or HL647-Aβ40 (red). The time axis is normalized to the half time of A*β*40wt. C-H: Representative TEM (C,D,E) and AFM (F,G,H) images of the fibrils after 24 h of incubation. The scale bars are 500 nm and 200 nm for TEM and AFM, respectively.

## MATERIALS AND METHODS

### Samples

Wild-type A*β*(1-40)-peptide (DAEFRHDSGY EVHHQKLVFF AEDVGSNKGA IIGLMVGGVV, A*β*40wt), its N-terminus cysteine mutant C-A*β*40, and the Atto488-A*β*40 peptide, labeled with Atto 488 (ATTO-TEC GmbH, Germany) were synthesized by the Core Unit of Peptide-Technologies of the University Leipzig. The peptides A*β*40wt and C-A*β*40 have been further labeled with fluorophores Atto 647N and Atto 655 (both from ATTO-TEC GmbH, Germany), in our laboratories. The labeled peptides HL488-A*β*40 and HL647-A*β*40 have been purchased from Anaspec, Inc., USA. To obtain different degrees of labeling (DOL), labeled and unlabeled A*β*40 peptides have been mixed with specific ratios. For example, for measurements with a 5% DOL the samples were mixed with A*β*40wt by a ratio of 1:20.

### Fluorescence labeling

#### Atto647N

1 mg/mL of A*β*40wt was dissolved in 25 mM sodium phosphate buffer (pH 8.3) and 17.5 mg/mL Atto647N NHS-ester (ATTO-TEC, Germany) in pure DMSO (Sigma Aldrich, Germany). Both solutions were first mixed and then incubated for 90 min in the dark at room temperature. Labeled A*β*40 was separated from the unbound free dye using a PD Minitrap G-25 column (GE Healthcare, U.S.A.). The final yield of labeled proteins was of about 2%.

#### Atto655

1 mg/mL of C-A*β*40 was dissolved in phosphate-buffer at pH 9.2 in the presence of 1 mM TCEP (Sigma Aldrich, Germany) and dialyzed in vacuum against the same buffer but at pH 7.5 and without TCEP. Atto655 maleimide (ATTO-TEC, Germany) was added at a 1.3:1 molar ratio and the solution stored for 2 h in the dark at room temperature. Labeled A*β*40 was separated from the unbound free dye using a PD Minitrap G-25 column (GE Healthcare, U.S.A.). The final yield of the labeled protein was 0.5%.

### Oligomerization and fibrillation

Following an established protocol (35), A*β*40-peptides were first dissolved in 25 mM sodium phosphate buffer (pH 9.2) with 150 mM NaCl. All fluorescently labeled samples were monomeric under these conditions as tested by Fluorescence Correlation Spectroscopy (see Fig. S1, Supplemental Material). Before fibrillation, A*β*40-samples were either dialyzed against the same buffer with a pH of 7.5 (4 h, MWCO 1000) or diluted by the same buffer of lower pH to reach pH 7.5. The final protein concentration was determined by absorption measurements and adjusted to 40 μM, if not otherwise stated. For fibrillation, A*β*40 solutions of 40 μM peptide concentration were incubated at 37 °C and shaken with 400 rpm - 450 rpm for 24 h with different sampling times. The appropriate incubation times to monitor oligomers or fibrils were chosen by TEM imaging, which, for A*β*40wt, revealed aggregates but no fibrillar structures after 4 h of incubation and mature fibrils after 24 h (see Fig. S2, Supplemental Material).

### Thioflavin T assays

A 2.5 mM Thioflavin T (ThT, Sigma Aldrich, Germany) stock solution prepared in 25 mM sodium phosphate buffer pH 7.5 containing 150 mM NaCl was stored at 4°C in the dark. Directly before the measurements it was filtered through 0.2 μm filters and diluted 1:10 in the same buffer. Monomeric A*β*40 solution in 25 mM sodium phosphate buffer pH 7.5 containing 150 mM NaCl was mixed with the ThT solution to a final ThT concentration of 20 μM. The samples were incubated in a FLUOstar omega well plate reader (BMG Labtech, Germany) at 37°C and 400 rpm with measurements at an excitation wavelength of 450 nm and the fluorescence emission detected at 480 nm, every 300 s.

### Transmission electron microscopy

5 μl of A*β*40-solution were dropped on Formvar/Cu grids with mesh 200 (Ted Pella, U.S.A.). After 3 min of waiting time the grids were first cleaned in water for 60 s and then negatively stained with 1% (w/v) uranyl acetate for other 60 s. TEM images were taken with an EM 900 (Zeiss) at 80 kV acceleration voltage.

### Atomic Force Microscopy

10 μl of fibrillated A*β*-solution were pipetted on top of freshly cleaved mica (Ted Pella, V1-grade, 10 mm diameter). After 1 min waiting time to allow sedimentation of the heavier fibrillar structures it was gently rinsed with clean water and dried in air. AFM measurements were taken with a Multimode 8 AFM (Bruker, U.S.A.) with NSG30 cantilevers (NT-MDT Spectrum Instruments Ltd., Ireland) in net-repulsive tapping mode choosing a target amplitude of 500 mV and a 5% peak offset.

### X-ray diffraction

The fibrillated dispersions of A*β*40wt or the fluorescently labeled peptides were placed in ultra-centrifugation at 60000 rpm for 1 hour for fibril sedimentation. The so obtained pellets were then either transferred into a ring-shaped aluminum holder (2 mm thick and with a central hole of 1.5 mm diameter) or scraped out with standard capillaries from Hilgenberg GmbH of borosilicate glass, with 3.8 mm or 1 mm outer diameter and 0.005 mm or 0.001 mm thickness, and left to dry overnight before measuring. Small and wide angle X-ray diffraction experiments (SAXS, WAXS) have being performed in transmission mode using a SAXSLAB laboratory setup (Retro-F) equipped with an AXO micro-focus X-ray source with an AXO multilayer X-ray optic (ASTIX) as monochromator for Cu-Kα radiation (λ = 0.154 nm). A DECTRIS PILATUS3 R 300K detector was used to record the two-dimensional scattering patterns. The measurements were performed at room temperature in vacuum over five samples all mounted on top of a grating with numbered positions used as collective holder. For those stored in glass capillaries, a measurement of the empty capillaries was taken for each q-range (*q_SAXS_* = 0.01 1.0 Å^-1^ and *q_W AXS_* = 0.4 3.2 Å^-1^) for background subtraction. Each intensity has been multiplied by a transmission factor to correct for absorption.

### Single-molecule Fluorescence Spectroscopy

#### Sample preparation

All single-molecule experiments were performed in 25 mM sodium phosphate with 150 mM NaCl at pH 7.5, unless otherwise stated. For single-molecule fluorescence spectroscopy 5 μl of the A*β*40-solutions were diluted to peptide concentrations of approx. 0.8 nM (DOL 5%) or 80 pM (DOL 100%), depending on the degree of labeling. After dilution, the samples were placed on top of a cover-slip for measurements. In case of an initial intensity loss due to adsorption of the peptide to the glass, the droplet was repeatedly renewed until no intensity change was observed within the first 5 minutes. The typical measurement time was 1 hour.

#### Measurement setup

Fluorescence measurements were conducted on a confocal microscope capable to perform polarization and time-dependent fluorescence measurements. The home-built optical microscope equipped with a tunable fiber laser (TVIS, Toptica, Germany, 1 ps pulse width, 80 MHz pulse frequency) and a diode laser (640 nm, LDH Series, Picoquant GmbH, 90 ps pulse width), a single mode fiber (SuperK FD7, NKPhotonics GmbH, Denmark) and a 60X/1.20 W PlanApo objective (Zeiss, Germany) is described elsewhere (38). The sampled fluorescence was collected and splitted into a parallel and a perpendicular polarization component by a broadband polarization-dependent beam-splitter (Melles Griot, U.S.A.) and focused onto two single-photon avalanche diodes (SPCM-AQR-14, Excelitas, Canada). Photon arrival time were recorded by a single photon counting board with 25 ps time resolution (TimeHarp 260 nano, PicoQuant GmbH, Germany).

#### Single-molecule data analysis

Intensity time traces were analyzed by selecting photons of either detection channel by a threshold criteria of 0.2 photons/ms and combining them into single bursts. Only bursts containing more than twice the average number of photons as detected for monomeric solutions were used to characterize oligomer solutions. To adjust for quenching effects, a fluorescence lifetime-corrected photon number, *N_phot_*, was determined using the equation *N_phot_* = (*N–N_bg_*)(τ_*F*,*mon*_/τ_*F,burst*_), where τ_*F,mon*_ is the average fluorescence lifetime of the monomer and *N_bg_* the fluorescent background of the buffer, as also used e.g. by (29). The applied selection criteria were optimized to reliably select not only bright molecules but also less intense molecules with long dwell times in the focus volume. After the determination of the encounter rates for each oligomer species, characterized by a certain relative photon number 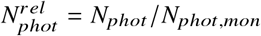, the apparent concentrations, *c*, were calculated by the equation 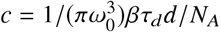, where *ω*_0_ is the width of the focus volume, *β* the encounter rate, τ_*d*_ the average diffusion time of all oligomers belonging to one oligomer species, *d* the dilution and *N_A_* the Avogadro number (25). For more details we refer to section S3 in the Supplemental Material.

#### Time-dependent fluorescence decay and anisotropy

Time-dependent fluorescence anisotropy experiments are used to measure the time-dependent depolarization of the fluorescence light due rotation of the emission dipole moment of the dye and can reveal specific fluorophore-peptide interactions. The time window of the experiment is the picosecond to nanosecond range, limited at short times by the pulse widths of the laser and the time response of the detectors (approx. 300 ps) and at long times by the fluorescence lifetimes. In case of unhindered rotation, the fluorescence anisotropy decays by a single exponential. As investigated in this study, the characteristic time constants for freely diffusive fluorophores, 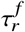, were between 450 ns and 700 ns (well below 1 ns) and the differences are related to the different sizes and shapes of the fluorophores. Fluorophores with attractive peptide interactions show a slower and potentially more complex decay. The rotation of the dye is either slowed down (fast binding and unbinding to the peptide) or the dye molecules adsorbs to the peptide which leads to a second decay component, 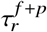, which can be related to the hydrodynamic radius *R_h_* of the peptide-fluorophore complex following the Einstein-Smoluchowski equation of the rotational diffusion with 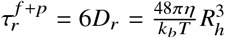, where *D_r_* is the rotational diffusion coefficient, *η* the solvent viscosity, *k_b_* the Boltzmann constant and *T* the temperature. In order to identify both processes, the normalized anisotropy was fitted using a two component exponential tail-fit: 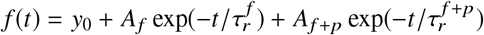. The ratio of the amplitudes *A_f_* _+*p*_/(*A_f_* + *A_f_* _+*p*_) can be taken as a measure for the attractive interaction strength. The offset, *y*_0_, which reflects the presence of few large aggregates which rotate on time scales longer than the fluorescence lifetime was subtracted if present.

## RESULTS AND DISCUSSION

In the following, the experiments which were conducted to reveal the effects of N-terminus functionalization of A*β*40-peptide with various fluorescent tags will be described and discussed in terms of (i) fibrillation kinetics, (ii) fibril morphology, (iii) LMW and HMW oligomeric species and (iv) fibrillar structure.

### Fibrillation kinetics

Two fundamental criteria to verify successful amyloid aggregation in time are the increase of *β*-sheet-rich structures and the final formation of bundles of extended *β*-sheets called fibrils. A common way to verify the *β*-sheet content of a sample and to follow the kinetics of *β*-sheet formation is to monitor the increase of the fluorescence of ThT. ThT is a small dye which binds to *β*-sheet-rich structures and shows enhanced fluorescence upon binding. Fig. 1B displays the time-dependent increase of the ThT fluorescence in the presence of A*β*40wt without and with 5% of the fluorescently labeled peptides Atto488-A*β*40 or HL647-A*β*40. We found, within the accuracy of the measurement, no major differences for the lag times and the half times, *T*_1/2_, which are the characteristic times of the onset time of the rise of the fluorescence and the time at which 50% of the maximum of the ThT fluorescence is reached. This result is in agreement with other literature findings, e.g. (28).

### Fibril morphologies

In order to investigate the final morphologies of the mature fibrils we used TEM and AFM imaging. The Fig.1(C-H) show representative images for A*β*40wt and two of the labeled peptides (more images can be found in Fig. S4 of the Supplemental Material). The TEM image of A*β*40wt displays elongated and twisted fibrils. Detailed analysis of the AFM image (Fig. 1F) revealed a ribbon like structure of approximately 20 nm width and 5 nm height and with a twist at every 120 nm. All these structural properties are in agreement with the common features of amyloid fibrils found in literature (40, 41). Fibrils of A*β*40wt formed in the presence of 5% of labeled peptides show slightly different morphologies, ranging from long and straight, which is very similar to the wild-type form and was observed e.g. for Atto488A-*β*40 (Fig. 1D), to thin and highly curved for HL647-A*β*40 (Fig. 1E). Interestingly, the degree of labeling (DOL) did not affect the final morphology. Even with a DOL of 100% Atto488-A*β*40 still formed elongated fibrils (see Fig. S4 of the Supplemental Material). Notably, the AFM image of HL647N-A*β*40 in Fig. 1H displays in addition to the fibrils also unstructured and soft material, whereas for Atto488-A*β*40 and A*β*40wt only few spherical structures are seen (Fig. 1G).

### Oligomer size distributions

If the concept of nucleation and growth is applied to the process of fibrillation, the formation of nuclei is the rate-limiting step of aggregation (42). Oligomers below a critical size are either thermodynamically instable (concept of one-step nucleation), or are stable but lacking in structure (concept of multi-step nucleation). However, if the size and/or structure of the critical nucleus is reached, larger aggregates will form rapidly. With increasing time all amyloid oligomers should participate in the fibrillar growth and finally disappear. In terms of oligomer size distributions this signifies that at short incubation times at least three populations should be found: i) a fraction of monomeric peptides, ii) a smaller fraction of (in)stable oligomers and iii) one of rapidly aggregating larger oligomers. Towards longer incubation times, one would expect a larger fraction of pre-fibrillar to fibrillar aggregates and a small fraction of monomers that did not aggregate. Moreover, aggregates which did not participate in the fibrillar growth due to lacking structure might also continue to grow with time and form large but unstructured oligomers. This will be fraction number iv)

To this end, time-dependent intensity traces of highly diluted A*β*40-samples with different fluorophores attached, were acquired at 0 h, and after 4 h and 24 h of incubation time. TEM imaging revealed for A*β*40wt after 4 h only spherical and prefibrillar oligomers only (see Fig. S2), which is in agreement with an estimated lag time of about 10 h, as determined by ThT-assays under identical conditions (35). Single-molecule experiments revealed in the time window of 2 to 6 h oligomer distributions, which did only slightly change, indicative of a quasi-equilibrium status between monomers and oligomers. Hence we chose in the following for all samples a constant incubation time of 4 h to investigate and compare the effect of fluorescence markers on oligomer size distributions.

Fig. 2 A and D displays the oligomer distributions as their concentration in function of monomer normalized number of photons as found for Atto488-A*β*40 and HL647-A*β*40-peptides, respectively. The initial monomer peptide concentration was 40 μM. The two distributions deviate strongly at any of the investigated sampling times: While Atto488-A*β*40 does not form HMW oligomers, after 4 h of incubation 20% of the oligomers of HL647-A*β*40-peptides are HMW oligomers. A reduction of the DOL to 5% allowed to measure at higher peptide concentrations, which enhanced the probability to detect HMW aggregates (Fig. 2B and E). Under this condition the dye-dependent variations were even stronger: while 5% of the oligomers of Atto488-A*β*40 are HMW oligomers, 59% of the oligomers of HL647-A*β*40 belong to this category. Moreover, the probability of the detection of very large aggregates with 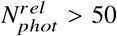 increases for HL647, which is in clear contrast to Atto488-A*β*40-peptides. Notably, the fluorescence properties of HL647 appeared to be more sensitive to environmental changes: while, monomers and oligomers of Atto488-A*β*40 have similar fluorescence lifetimes of 3.85 ns (Fig. 2C), the fluorescence lifetime of HL647-A*β*40-peptides increases if oligomers are formed. Fibrils, however, show a strongly reduced fluorescence lifetime of 1.3 ns. In Fig. 2 F the normalized distribution is shown for three different environments. The monomeric peptides were fist dissolved in buffer at pH 9.2. After changing the pH to 7.5, the majority of molecules displayed a small shift of the fluorescence lifetime from 1.3 ns to 2.0 ns, which can be explained by a weak pH-dependence of the emission properties of the fluorophore itself. But, in addition, aggregation was initiated and some oligomers were instantaneously formed. The formed oligomers displayed an average fluorescence lifetime of 2.8 ns. Hence, the detected photon numbers did not only change due to the increase in the number of aggregated peptides but also by a shift of τ_*F*_. This behavior indicates an increased solvent protection of HL647-molecules and reveals the hydrophobic nature of the fluorophore. The relative photon numbers used in the size distributions were corrected for fluorescence quenching (see Material and Methods section). Without this correction, lower quenching could be misinterpreted by an apparent increase in size. Interestingly, the fibrils formed by HL647-A*β*40-peptides displayed a highly quenched fluorescence lifetime of approx. 1.0 ns.

**Figure 2:**
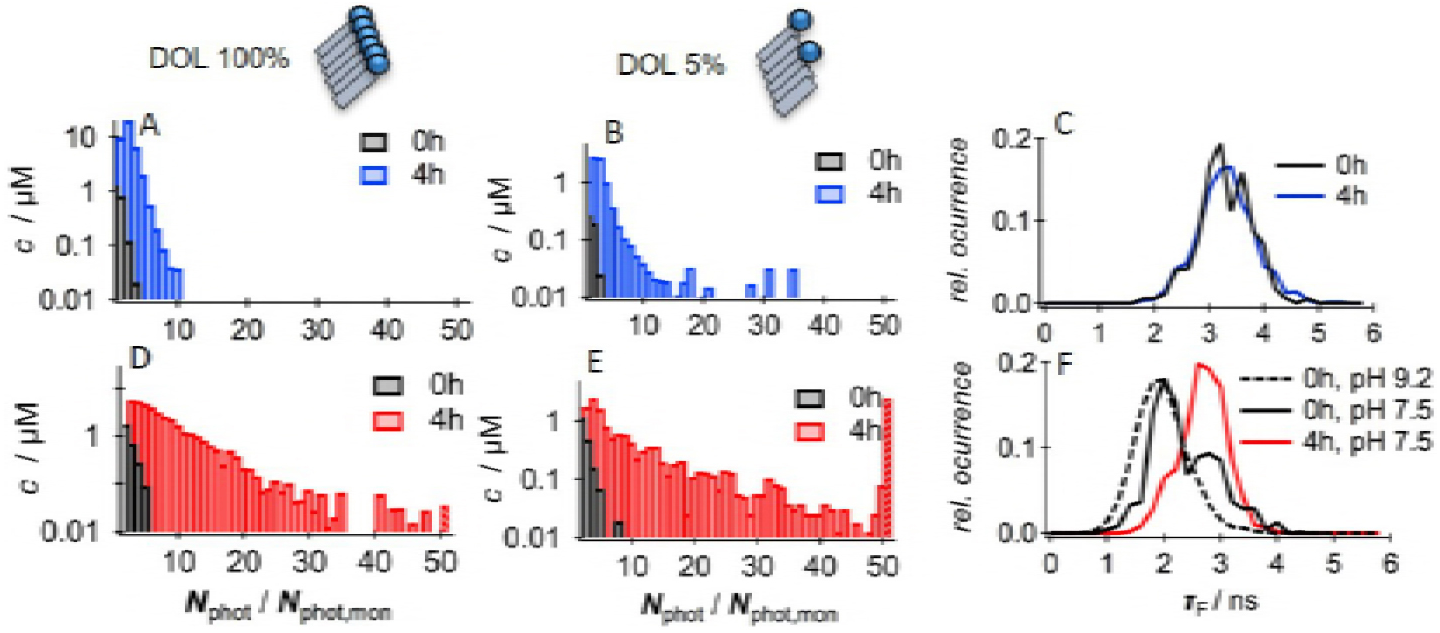
The effect of the DOL of N-terminus bound fluorophores on oligomer size distributions of A*β*40. A,B,D,E): Concentration distribution of oligomers in terms of relative photon numbers *N_phot_/N_phot,mon_* using 100% (A,D) and 5% (B,E) of labeled A*β*40-peptides.The oligomer distribution are shown before (black) and after 4 h of incubation (colored) for Atto488-A*β*40 (blue) and HL647-A*β*40 (red). The last bar in the histograms sums up all oligomers above 50x*N_phot_/N_phot,mon_*. C,F: The distributions of the fluorescence lifetimes, τ*_F_*, for all detected molecules before (black) and after 4 h of incubation (colored) for Atto488 (blue) and HL647 (red). The dashed line in F displays HL647-Aβ40 dissolved in pH 9.2, all other measurements were conducted at pH 7.5 and an initial peptide concentration of 40 μM. The selection criteria were chosen such that the probability to include monomers in the histograms is very low.

In the last section the question should be addressed, how the discussed oligomer distributions can be related to size. The relative photon numbers used to characterize the oligomers depend not only on the number of fluorescent labels, which should be proportional to the aggregation number, but also on the individual dwell times in the focus volume. Hence, any increase in the number of fluorophores as well as the reduction of the translational diffusion coefficient will lead to enhanced photon counts. However, the unexpectedly narrow distribution of the molecular brightness led to an apparent linear relationship between the average diffusion time, τ_*d*_, and 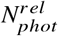 which justifies to relate 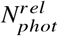 to size: 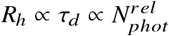 with 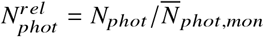 (see Fig. S3 for more details).

### High molecular weight oligomers

In the following, the effect of five different fluorophores were investigated. In order to pronounce HMW oligomers, which we call oligomers with 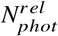 larger than 10, a low DOL was used. Fig. 3 displays the result and links characteristic features in terms of oligomer size distributions, the shift of fluorescence lifetimes upon aggregation and the time-resolved fluorescence anisotropy of monomeric peptides. Starting with the oligomer size distributions, we found for Atto488-A*β*40-peptides a narrow distribution which can be characterized by an average value of *N*̅*_phot_* = 2.7 *N*̅*_phot,mon_* and a standard deviation of 4.7 *N*̅*_phot,mon_*. Just very rarely, some large aggregates with *N_rel_* > 50 were detected. After 24 h of incubation the detected oligomer sizes were again strongly reduced and comparable to the distribution before fibrillation (see also Fig. 2B). Oligomers of HL488-A*β*40-peptides had a slightly increased average value of 4.1 *N*̅ *_phot,mon_*. The distribution width was 6.3 *N*̅*_phot,mon_*, but like for Atto488, most of the oligomers were consumed by the end of the fibrillation process. In comparison, Atto655-A*β*40 formed larger oligomers with an average of 7.4 *N*̅*_phot,mon_*. The standard deviation of the distribution was 8.8*N*̅*_phot,mon_*. Atto647N-A*β*40 and HL647-A*β*40 display a clear shift to even larger aggregates. The average values of the distribution were 10.9 and 10.6 *N*̅*_phot,mon_* and the standard deviations 14.7 and 14.9 *N*̅*_phot,mon_*, respectively. Very large aggregates of above 50 *N*̅*_phot,mon_* were frequently detected but not included in the average and standard deviation of the distribution. It can be assumed that these large aggregates are no oligomers in the narrow sense. Notably, the larger aggregates maintained in the solution, even after 24 h of incubation, whereas the fraction of smaller aggregates seemed to be reduced. In Fig. 3C the fluorescence lifetimes of the detected bursts before and after 4 h of incubation for all investigated fluorophores are compared. It appears that all dyes which formed HMW aggregates, thus Atto655, Atto647N and HL647, displayed a clear shift to higher fluorescence lifetimes if aggregates were formed. We refer this shift to hydrophobic properties of the dyes and to solvent-shielding upon aggregation. In contrast, Atto488 and HL488 do not show any difference, which reveals a more hydrophilic nature and indicates, that Atto488 and HL488 stay exposed to the solvent during fibrillation.

**Figure 3:**
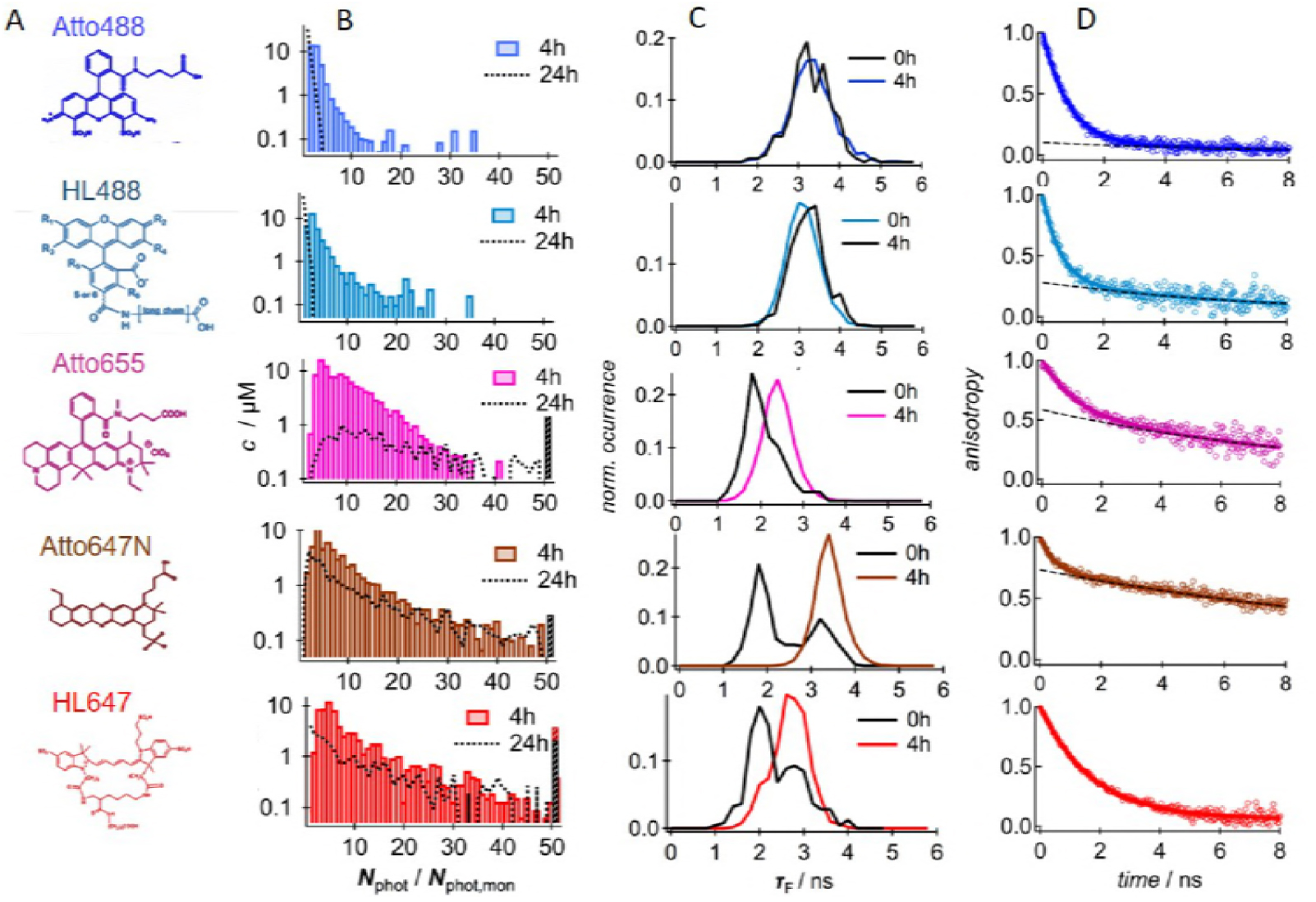
The effect of N-terminus bound fluorophores on oligomer size distributions of A*β*40 at low DOL. Chemical structures (A), concentration distribution of oligomers in terms of relative photon numbers *N_phot_/N_phot,mon_* of labeled A*β*40-peptides (A655-A*β*40 0.5%, A647N-A*β*40 2%, all others DOL5%) after 4 h and 24 h (B), the fluorescence lifetime distribution before and after 4 h of fibrillation and the time-dependent anisotropy of the fluorescent labels before fibrillation for 5 different fluorophores.

The normalized time-dependent anisotropy measurements of Fig. 3D display the time-dependent depolarization of the fluorescence emission of the dyes, which can be due either to fast orientation fluctuations of the dipolar moment of the dye abouts its own axis (fast component) or to the rotation of the dye-peptide complex (slow component). For Atto488-A*β*40-peptides the majority display a fast decay with τ_*r*_ =(510±10)ps. However, there is also a small fraction of 12% with have a longer decay time of of (7.00±0.18) ns which refers to a hydrodynamic radius, *R_h_*, of approximately 1.9 nm. This value is slightly larger than the radius of 1.6-1.7 nm found for unlabeled Ab40-peptides (43, 44). HL488-A*β*40-peptides show a similar behavior: in addition to a fast anisotropy decay of unhindered rotation, 28% of the fluorophores show a decay on the order of the fluorophore-peptide complex. Atto655 and Atto647N show again two different decay times, but the fraction of fluorophore-peptide complexes is much larger as compared to Atto488 and HL488. 62% of Atto655-A*β*40 and 74% of Atto657N-A*β*40 display a slow decay with (6.08±0.05) ns and (6.48±0.05) ns for Atto647N and Atto655, respectively. We assume that the enhanced interaction is mediated by the increased hydrophobicity of the dyes. In contrast to all other dyes, HL647-A*β*40-peptides display a single decay only, which might be related to its different chemical structure. However, the decay has a time constant of τ_*r*_=(1.77±0.02) ns, which is apparently larger than the decay of (0.89±0.05) ns observed for peptides at elevated pH (pH 9.2). Hence, at pH 7.5 all fluorophores strongly interact with the A*β*40-peptides, but do not form static complexes.

### Fibrillar structure

In order to investigate whether the enhanced interaction of fluorophores with the peptide has any impact to the fibrillar structure, X-ray scattering experiments were conducted. X-ray diffraction can be used for determining the atomic or molecular structure of crystals or crystal-like samples. Relative distances and absolute positions of the first neighboring atoms can be deduced from the scattering intensity and scattering directions of the X-rays deviated by the atomic electronic densities. The first X-ray diffraction measurements of A*β* amyloid fibrils trace back to the latest ’60s (45, 46) and revealed a typical cross-*β* structure with *β*-sheets arranged in parallel to the fibril axis and their constituent *β*-strands perpendicular to the fibril axis. The latter resulted in 4.7 to 4.8 Å meridional reflections and a 10 Å equatorial reflection respectively, which are considered the hallmarks of a cross-*β* structure (47). All amyloid proteins or peptides show indeed very similar X-ray diffraction patterns (40). In Fig. 4 just the meaningful WAXS region between q=0.4 Å^-1^ and 1.8 Å^-^1 of the curves is reported, after amorphous background subtraction and normalization with respect to the areas below the second peak, which indicate the crystalline content due to *β*-sheets formation. The full accessible *q*-range is shown in Fig. S5 (Supplemental Material). The vertical lines in the Fig. 4 indicate peaks positions of A*β*40wt which are related to the *β*-strand distances along the fibril axis and the *β*-sheet distance perpendicular to it. The first refers to the peak at *q* = 1.32 Å^-1^, indicating a distance of 4.75 Å between β-strands, and the latter to the peak at 0.58 Å^-1^, which is related to a distance of 10.8 Å between *β*-sheets. The equatorial reflections appear broader and weaker than their meridional equivalents implying a much lower crystalline order in directions perpendicular to the fiber axis then parallel to it. The scattering intensity of the labeled peptides display the same two peaks with q-values ranging from 1.32 Å^-1^ to 1.34 Å^-1^ for the meridional reflections and from 0.63 Å^-1^ to 0.65 Å^-1^ for the equatorial ones, corresponding to distances ranging between 4.7-4.75 Å and 9.6-10.8 Å respectively. More precisely, all the HiLyte labeled samples (HL488-A*β*40, HL647-A*β*40) show a peak maximum which refers to a distance of about 10.0 Å, which is significantly lower compared to A*β*40wt. To compare Atto488-A*β*40, this peptide shows at a low DOL of 5% no difference from A*β*40wt, but if the DOL is increased to 100%, a shift to even higher q-values with an inter-*β*-sheet distance of 9.6 Å. However, the common reflection for the distances of the *β*-strands (4.75 Å) indicates very similar, if not identical, nuclei. With other words, the fluorophores did not affect the structure of the nuclei, as this structure would propagate during growth. However, the lateral order of the fibrillar bundles was modified, which is depicted by the dye-dependent differences of the other peak.

**Figure 4:**
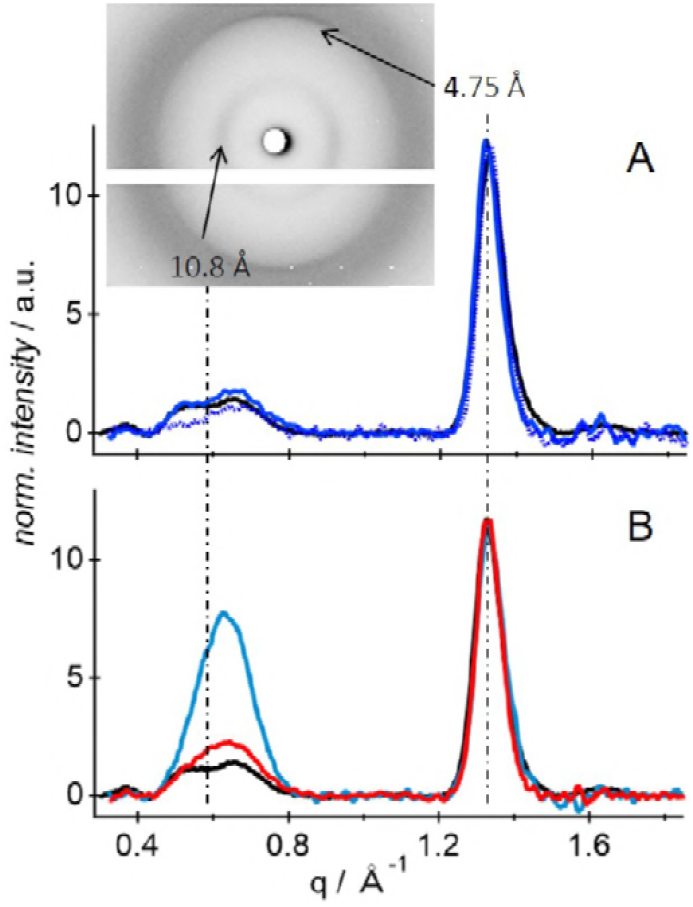
The effect of N-terminus bound fluorophores on the fibrillar structure of A*β*40. Relative scattering intensity of the amyloid fibrils in the range of 0.4 and 1.8 Å^−1^ after azimuthal integration, background subtraction and renormalization with respect to the crystalline *β*-sheet content. A: A*β*40wt (black line), Atto488-A*β*40 (blue solid line for DOL 5% and blue dashed line for DOL 100%). B: A*β*40wt (black line), HL488-A*β*40 (blue line) and HL647-A*β*40 (red line), both DOL 5% Inset: 2D image of Atto488-A*β*40 (DOL 5%) with the typical meridional and equatorial peaks reflections at 4.75 Å and 10.8 Å respectively.

## SUMMARY

In this study the impact of N-terminal labeling of A*β*40-peptides was investigated using different common fluorophores with different spectral properties and hydrophobicity. In the following section the experimental findings for each dyes are summarized and briefly discussed.

### Atto488-A*β*40

Fluorescence anisotropy measurements revealed that most of the bound Atto488 fluorophores display unhindered rotational motion and only a low percentage of 12% of the dye molecules show cooperative rotation with the peptide. This finding correlates with the successful formation of elongated, mature fibrils which are very similar to the morphology of A*β*40wt-fibrils as found by TEM and AFM imaging. X-ray diffraction reveals inter-strand and inter-*β*-sheet distances identical to A*β*40wt. Atto488-A*β*40 forms only small oligomers as found by single-molecule fluorescence experiments, which were consumed during fibrillation. Hence Atto488-A*β*40 is an excellent candidate to study oligomerization and fibrillation.

### HL488-A*β*4 and HL647-A*β*40

HL488-A*β*40 and HL647-A*β*40 are two peptides which are commercially available and therefore often used in literature, e.g.(19, 29–31, 36). A clear drawback of the use of these dyes is their pH-dependence in terms of net-attractive interactions with the A*β*40-peptide itself, which was discussed for HL647, but could been observed somewhat weaker for HL488 as well. At a pH of 7.5 23% of the HL488 dyes and basically all of HL647 show slower rotational motion compared to pH 9.2. Moreover, HL647-A*β*40 formed large oligomers and promoted the formation of very large aggregates with 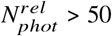, whereas for HL488-A*β*40 mainly small oligomers with an average value of 4xN̅*_phot,mon_* were detected. The latter is in agreement with the small oligomers of up to 6 monomers, which were found, e.g. by photo-bleaching experiments of surface attached HL488-A*β*40-peptides (19). The tendency of HL647-A*β*40 to form small fractions of large aggregates at the early stage of fibrillation is mentioned in literature as a side note (30). Interestingly, the differences in terms of interaction and oligomer sizes can be also linked to a different morphology of the fibrils after 24 h of fibrillation. While HL488-A*β*40 forms straight and extended fibrils similar to Atto488-A*β*40, the fibrils of HL647-A*β*40 seem to be thin and strongly bended. Delayed fibrillation kinetics cannot be the reason for the different morphologies as standard ThT-assays used to investigate the kinetics of *β*-sheet formation display no delay for HL647-A*β*40 compared to A*β*40wt. And since the local structure of the fibril as investigated by WAXS measurements do not show any deviation from the wild-type as well, we conclude that for HL647-A*β*40 not the nucleation but the fibril growth is modified, e.g. by termination of the growth end and hindered lateral stacking.

### Atto655-A*β*40,Atto647N-A*β*40

Atto655 and Atto647N are examples for a hydrophilic and a hydrophobic dye in the red spectral range. Surprisingly, both fluorophores tend to stick to the peptide in the buffer conditions used, however, oligomer size distributions show for the more hydrophobic fluorophore Atto647N a broader distribution. This corresponds to literature finding which report on the use of Atto647N the formation of high MW oligomers, with e.g. a 70-fold increase of the diffusion time (27) or a time-independent, 10-fold increase in the diffusion time (48). The latter value agrees well with our finding of *N*̅_*rel*_ of 10.9. Notably, both peptides form shorter and thinner fibrils compared to A*β*40wt or Atto488-A*β*40.

## CONCLUSION

Combining the results discussed above, we conclude that the formation of amyloid oligomers is very sensitive to the chosen fluorophore, while the fibrillation kinetics and the structures of the final fibrils are only marginally affected. In particular, the occurrence of high MW oligomers of N-terminally labeled A*β*40-peptides can be linked to hydrophobic properties of the attached dye, which is indirectly pictured by i) attractive fluorophore-peptide interactions and ii) by reduced fluorescence quenching upon aggregation. Since high MW oligomers tend to maintain in the solution after fibrillation, we further conclude that they do not contribute to the fibrillation. Consequently, low MW oligomers form the nucleus and control fibrillar growth, which explains the similar fibrillation kinetics of labeled peptides as well as the same internal structures along the axis of fibrillar growth (4.75 Å peak). Differences were found on larger scales, ranging from the distance between *β*-sheets (10 Å peak) to the overall morphology of the fibrils on the 100 nm-scale (TEM images). Consequently, the ideal fluorophore which should be used to investigate oligomeric states of A*β*40 should not lead to high MW oligomers, which would superimpose with those responsible for fibrillation. From our studies we found Atto488 and also HL488 as promising candidates for such kind of labeling functionalization, which may function for other kind of amyloid peptides as well.

## AUTHOR CONTRIBUTIONS

M.O. designed the research, J.W., S.D.S. and B.V. performed the research, J.W. and S.D.S. contributed equally. J.W.,S.D.-S. and M.O. analyzed the data. M.O., S.DS., J.W. and J.B. wrote the manuscript.

## ACKNOWLEDGMENTS

The authors would like to acknowledge Dr. Sven Rothemund (University Leipzig) for peptide synthesis, Monika Baumann (Martin-Luther-University Halle-Wittenberg) for initial TEM images and Prof.Thomas Thurn-Albrecht (Martin-Luther-University Halle-Wittenberg) for providing access to the SAXS equipment and fruitful discussion. The study was supported by a grant from the Deutsche Forschungsgemeinschaft (DFG TRR 102, B12).

## SUPPLEMENTARY MATERIAL

An on-line supplement to this article can be found by visiting BJ Online at http://www.biophysj.org.

